# DNA polymerase ζ is a robust reverse transcriptase

**DOI:** 10.1101/2024.09.27.615452

**Authors:** Ryan Mayle, William K. Holloman, Michael E. O’Donnell

## Abstract

Cell biology and genetic studies have demonstrated that DNA double strand break (DSB) repair can be performed using an RNA transcript that spans the site of the DNA break as a template for repair. This type of DSB repair requires a reverse transcriptase to convert an RNA sequence into DNA to facilitate repair of the break, rather than copying from a DNA template as in canonical DSB repair. Translesion synthesis (TLS) DNA polymerases (Pol) are often more promiscuous than DNA Pols, raising the notion that reverse transcription could be performed by a TLS Pol. Indeed, several studies have demonstrated that human Pol η has reverse transcriptase activity, while others have suggested that the yeast TLS Pol ζ is involved. Here, we purify all seven known nuclear DNA Pols of *Saccharomyces cerevisiae* and compare their reverse transcriptase activities. The comparison shows that Pol ζ far surpasses Pol η and all other DNA Pols in reverse transcriptase activity. We find that Pol ζ reverse transcriptase activity is not affected by RPA or RFC/PCNA and acts distributively to make DNA complementary to an RNA template strand. Consistent with prior *S. cerevisiae* studies performed *in vivo*, we propose that Pol ζ is the major DNA Pol that functions in the RNA templated DSB repair pathway.

## Introduction

DNA damage plagues all cell types. Damage comes in the form of any alteration in DNA structure leading to a change in coding capacity or deterioration of genome integrity. Different types of radiation, environmental toxins, and endogenous oxidation products and metabolites can introduce an enormous range of chemical changes in DNA structure. Self-induced damage can be incurred by collisions and aberrant action of endogenous protein complexes that replicate, transcribe, and alter the topology of DNA. To cope with such a vast array of damage, cells have an arsenal of countermeasures to restore genomic integrity. This includes an assortment of mechanical systems to traverse or remove damaged bases, restore tracts of inappropriate nucleotides, and rejoin broken DNA molecules. With certain notable exceptions of direct reversal of DNA damage by photoreactivation of ultraviolet light-induced cyclobutane dimers (1) and demethylation of alkylated bases via dioxygenases or suicide demethylases (2), a common feature of all DNA repair mechanisms is the requirement for some amount of DNA synthesis. This could range from incorporation of a mere few nucleotides in the case of excision repair of a single modified base (3), to hundreds of nucleotides in the case of DNA double-strand break (DSB) repair (4), or even thousands in the case of break induced replication (5).

Specialized DNA polymerases exist to repair short gaps created after base excision (3), traverse damaged nucleotides in concert with ongoing DNA replication (6,7), and to blunt or polish broken DNA ends in preparation for rejoining by non-homologous end joining (8). Repair of DSBs by the homologous recombination (HR) pathway requires longer tracts of synthesis, and is thought to utilize the replicative DNA Pol δ in a migrating D-loop reaction that provides part of the means for bridging the broken DNA ends (9). In HR the conventional template used for repair synthesis is an undamaged homologous DNA partner, usually the sister chromatid (10).

Evidence has emerged from recent studies suggesting that RNA transcripts might also be directly utilized as templates in DSB repair (11,12). In this case, DNA synthesis would arise from a reverse transcriptase (RT) activity. In studies of RNA templated repair performed in *Saccharomyces cerevisiae*, reverse transcriptase activity was in some cases attributed to that encoded by the Ty element retrotransposon (13). However, this is believed to function through a cDNA intermediate, where the RNA is only indirectly used as a repair template (14,15). Pol θ of mammalian cells has also been shown to harbor a robust reverse transcriptase activity (16) and is required for DSB repair by the microhomology mediated end joining pathway (17). Recent biochemical studies have shown that Pol η has some reverse transcriptase capability (18) and can promote RNA templated repair *in vivo* (19). New genetic investigations in yeast have suggested that Pol ζ performs the RT activity necessary for RNA templated repair (13). In contrast to the cDNA mediated Ty RT repair pathway, these newly identified repair pathways are proposed to involve repair synthesis directly templated by RNA. An example of such a repair mechanism is illustrated in Figure 1, where a transcript mRNA is used to bridge a DNA DSB and facilitate repair via RT synthesis.

**Figure 1.**
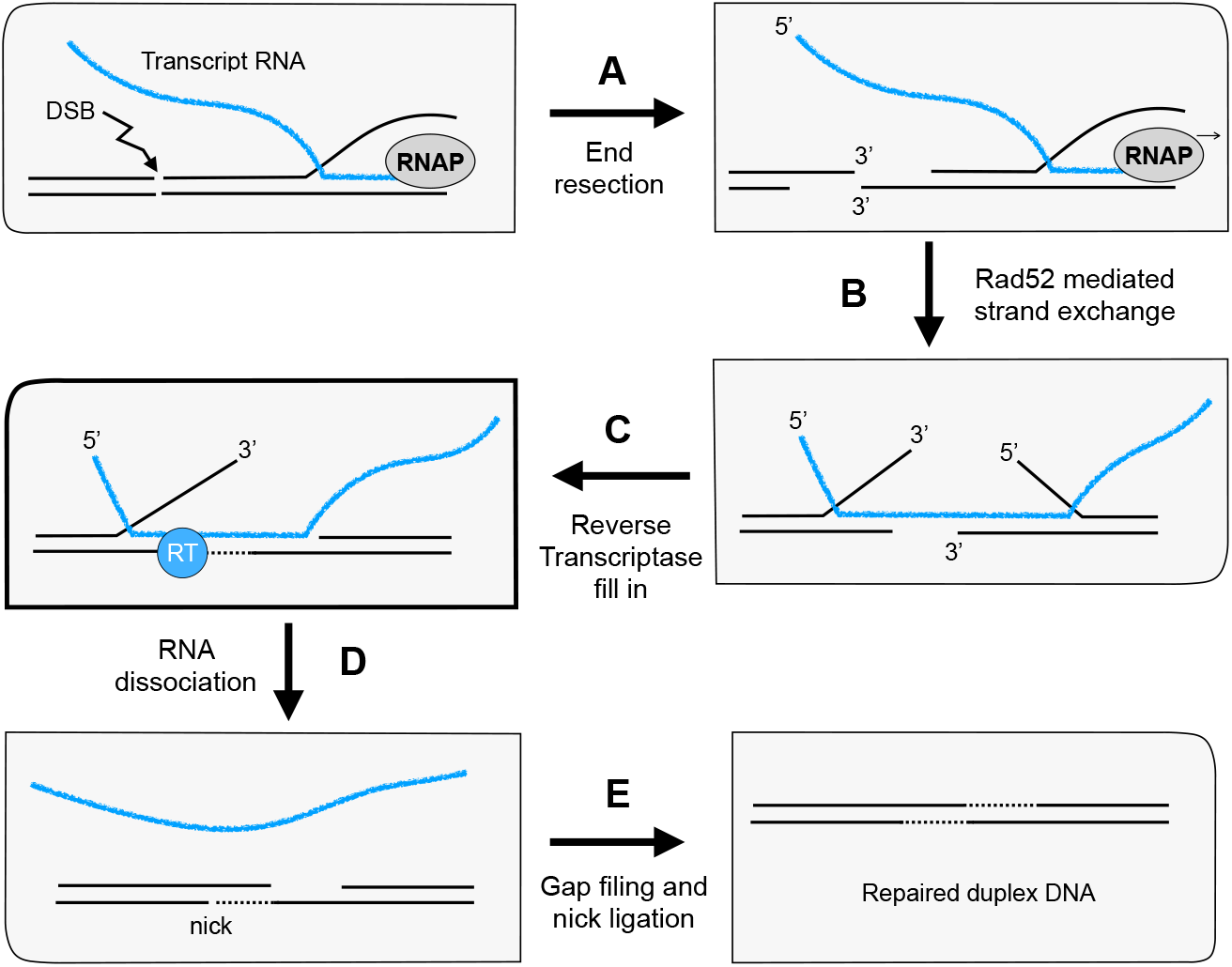
Model of RT activity for RNA mediated double strand DNA break repair. A dsDNA break may occur after transcription has produced an mRNA transcript. In this event, any resection and consequent loss of DNA information can be “rescued” by the mRNA transcript as follows. Following (**A)** end resection, (**B)** the mRNA can bridge the DSB via annealing and Rad52-mediated strand exchange. After this, (**C)** the gap created by resection can be filled in by a reverse transcriptase. After (**D)** the RNA dissociates from the break site, (**E)** normal DNA templated gap filling and nick ligation restore an intact duplex.

To help address the gap in knowledge about DNA Pol RT activity in this important field, we have purified all seven nuclear DNA Pols of budding yeast and compared their ability to perform reverse transcription. We find that Pol ζ contains the most proficient RT activity by far. Although comparison of extension by Pol ζ on RNA compared to DNA templates shows greater activity on DNA, Pol ζ’s RT activity is more robust than any of the other DNA Pols, and is not hindered by RPA, RFC, or PCNA. Thus, the results of this report suggest that Pol ζ may be the major enzyme involved in the RNA directed DSB repair pathway, consistent with cellular studies (13).

## Results

We have purified all seven of the nuclear DNA Pols of *Saccharomyces cerevisiae* (Fig S1) and have tested them for relative DNA polymerase and reverse transcriptase (RT) activity. These include the replicative polymerases Pol α, δ and ε, the translesion polymerases Pol η, ζ, and Rev1, and the X-family polymerase Pol IV, considered the equivalent of metazoan Pol β, which is dedicated to base excision repair (20). As shown in Fig. 2a, these DNA Pols can efficiently utilize dNTPs to extend a primer across a 30 nt ssDNA template strand, with the exception of Rev1 and Pol IV. Rev1 was previously characterized in many systems to be a very specialized Pol, and typically only incorporates a single dC nucleotide (21). The activity of Pol IV is less well characterized than other yeast polymerases, but it was shown to fill short gaps and act distributively to extend at longer single strand regions (22-24).

**Figure 2.**
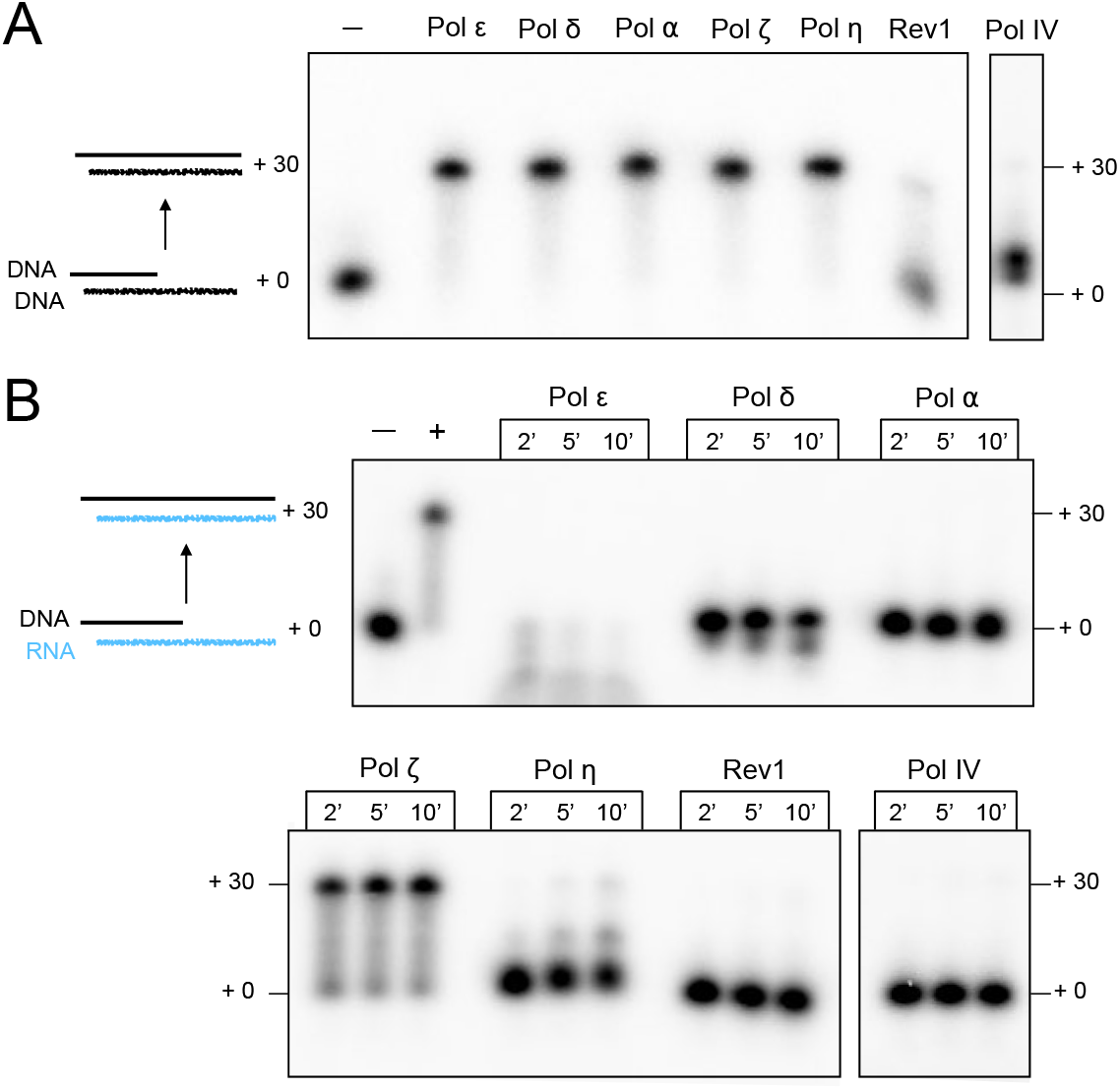
Pol ζ is the only yeast Pol having significant Reverse Transcriptase activity. **A)** comparative DNA synthesis activity using 20 nM of either Pol ε, δ, α, ζ, or η. Elevated concentrations were used for Rev1 (60 nM) and Pol IV (100 nM) in order to observe any extension activity. **B)** RT activity of each Pol, using the same fmol of Pol as in panel A.

This report tests these DNA Pols for RT compared to DNA polymerase activity using a primed substrate with comparable RNA or DNA template strands. The results show that all the Pol preparations have DNA Pol activity, although Rev1 shows minimal extension activity as expected (**Fig. 2a**). Upon examination of RT activity using an RNA template strand of the same sequence of the DNA template, we observe that only Pol ζ has significant RT activity (**Fig. 2b**). Note, however, that Pol η does have some RT activity, consistent with previous reports (18,25). However, at equal molar concentrations Pol η has substantially weaker RT activity than Pol ζ. Pol ε, and to a lesser extent Pol δ, degrades the primer rather than extending it when utilizing an RNA template strand. We do not rule out the possibility that Pol δ and Pol α may extend a nucleotide or two on RNA, as was previously reported (12).

Next, we asked whether Pol ζ RT activity is processive or distributive in the context of extending 30 nucleotides to the end of our ^32^P-DNA primed RNA substrate. To determine this, we titrated Pol ζ into reactions containing a constant amount of DNA primed RNA template. If Pol ζ were processive, the products would not show a dependence on the amount of Pol ζ added, but would show more of the same product at different concentrations. If Pol ζ were distributive, the product length would depend on the concentration of the polymerase added. The results (**Fig. 3**) show that product length depends on the concentration of Pol ζ, demonstrating that the reverse transcriptase activity of Pol ζ is distributive, and that processivity is less than 30 nucleotides in our assay conditions.

**Figure 3.**
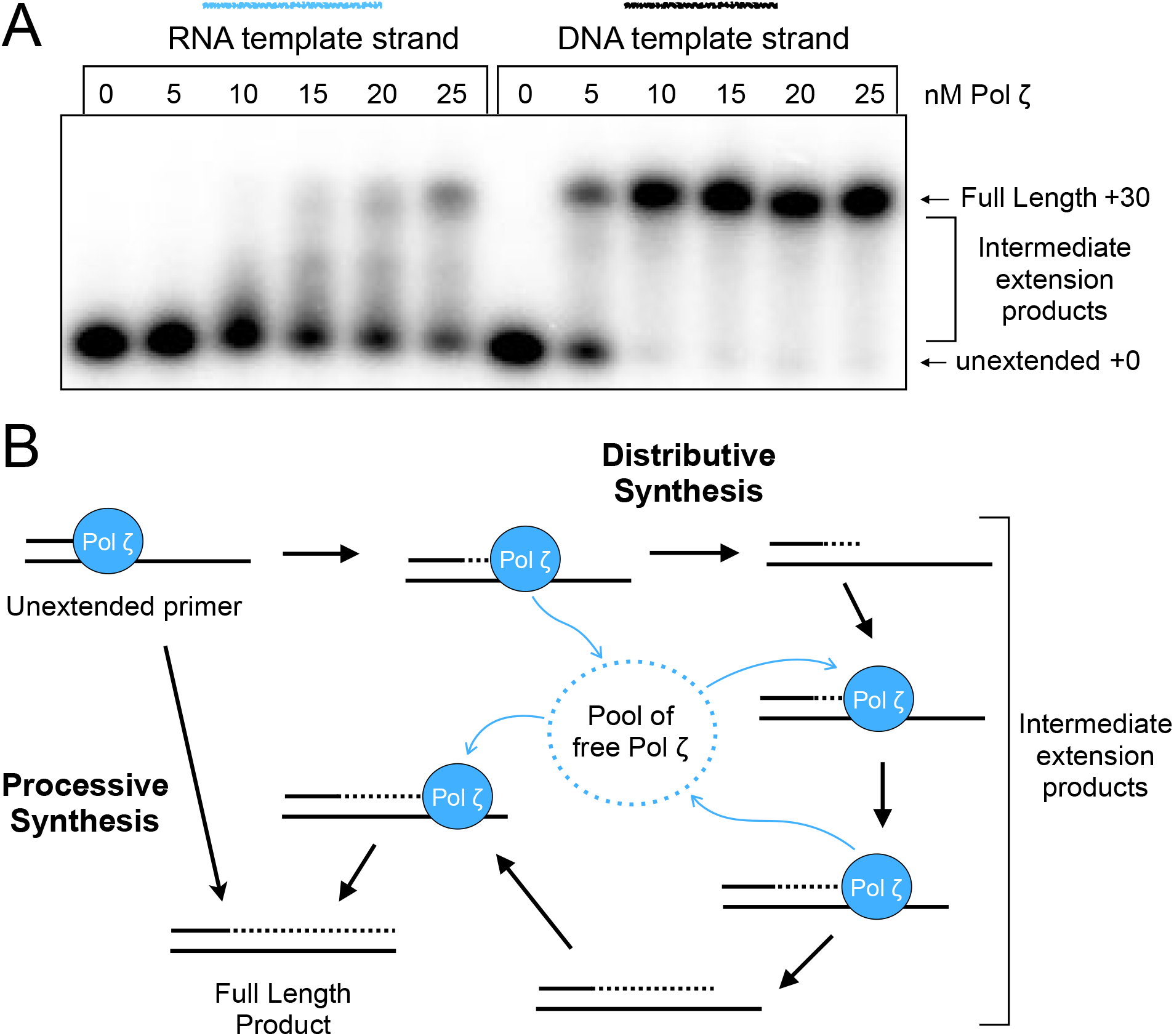
Pol ζ is distributive during RT activity. **A)** Titration of Pol ζ into extension assays with DNA primed RNA or DNA templates. RNA template reactions were stopped at 1 minute and DNA template reactions at 10 seconds. Products are analyzed in a 10% urea PAGE. **B)** Diagram describing the two possible modes of synthesis for Pol ζ, processive or distributive.

One may consider whether a distributive enzyme might be hindered by the single strand binding protein, RPA, which would undermine the view that Pol ζ might be a functional RT *in vivo*. To test this, we first examined binding of RPA to the RNA template primed with DNA under our primer extension assay conditions, by using an EMSA assay (**Figure S2**). We observed that RPA binds the DNA primed RNA template when present at 100 nM, which matches the concentration used for RT assays performed in this report (**Fig. S2**).

Addition of RPA in extension assays of Pol ζ RT activity does not result in a significant inhibitory nor stimulatory effect (**Fig. S3 A-B**). We also tested the effect of adding RFC, PCNA or RFC + PCNA to the Pol ζ RT assays (**Fig. S3 C-D, Fig S4A**). We observed no notable inhibition or stimulation of Pol ζ RT activity by RFC and PCNA when added individually, or when added together. This could be due to RFC not loading PCNA on a DNA primed RNA template, consistent with a prior investigation of RFC’s loading abilities (26). We also observed that RFC + PCNA does not stimulate Pol ζ on a primed DNA substrate (**Supplementary Fig. 4B**). Given these results, it may not be surprising that RFC + PCNA does not have an impact on RT activity.

## Discussion

There are several distinct pathways of DNA repair, as explained in the Introduction. Among these pathways is a mechanism for DSB repair that uses the sequence of a transcript mRNA to regain any lost DNA information through use of a reverse transcriptase. This can be accomplished by annealing of the RNA transcript across the DNA double strand break followed by DNA fill-in using the mRNA transcript as a template (**Figure 1**). The idea that Pol ζ might be the RT for RNA template directed repair was suggested by earlier genetic studies (13). While a yeast transposable element RT initially confused the issue, it was eventually shown that in its absence the chromosomal encoded genes that comprise Pol ζ were required for this method of DSB repair. Although the genetic studies suggested that Pol ζ might be the enzyme responsible for the reverse transcriptase activity, whether Pol ζ possessed RT activity had not been directly tested.

In this report, we purified each of the seven nuclear DNA Pols of *S. cerevisiae*. Upon examination of these different DNA Pols, we find that Pol ζ has by far the most robust RT activity. Interestingly, Pol η is reported to contribute RT synthesis relevant to DSB repair (18,19). However, the relative RT activity of human Pols η and ζ is not known, and thus it remains to be determined whether the results of this report, using budding yeast Pols, generalize to human Pols. Importantly, Pol ζ has not been extensively studied in human cells in the context of RNA templated repair and may possess similar activity to the budding yeast system explored here. Nonetheless, it is an exciting time for a deeper understanding of this newly discovered DSB repair pathway, and this report serves as a notable finding confirming the prediction of genetic studies that Pol ζ may possess RT activity (13). We hope to build on this work using recombination enzymes along with DNA Pol ζ and its associated machinery to reconstitute this transcription based DSB repair process.

## Experimental Procedures

### Proteins

Pol ε, Pol δ, Pol α, RFC, RPA and PCNA were purified as previously described (27). Each of Pol ζ, Pol η, Rev1, and Pol IV were cloned into the following integration vectors. Pol eta with an N-terminal 3X FLAG tag was cloned into pRS403 (His^+^ selection) under control of the Gal1 promotor. N-terminal 3X FLAG tagged Pol IV and Rev1 were each cloned into pRS404/GAL (Trp^+^ selection) under control of the Gal1 promotor. The four subunit Pol ζ (Rev3/Rev7/Pol31/Pol32) was cloned into three integration vectors: Rev3 with a C-terminal 3X FLAG tag was cloned into pRS402/GAL (Ade^+^) under the Gal1 promotor. Rev7 was cloned into pRS405/GAL (Leu^+^) under the Gal1 promotor. Pol31 and Pol32 were cloned into the pRS403/GAL (His^+^)) under the Gal1 and Gal10 promotors, respectively. The integration plasmids containing the genes of Pol η, Rev1, Pol IV and the 4-subunit Pols ζ were integrated into yeast strain OY01 (27). Unlike the other Pols, Pol ζ is four subunits and required integration of two plasmids instead of one, as noted above.

24L of each strain encoding either Pol ζ, Pol η, Pol IV was grown to 0.6 OD_600_ and induced with galactose for 6h at 30°C as previously described for Pols δ and α (28). The Rev1 strain was grown similarly, to 0.6 OD_600_, except was induced at 26°C after only 4h of Gal induction. The DNA Pols were purified similarly using the following protocol. 24L induced yeast cells were lysed in a SPEX cryogenic mill as described (27). Lysed cells were clarified by centrifugation and 1-2 ml anti-flag beads (Sigma, #A2220) were added to the supernatant (about 150 ml), followed by 1 h on a rocking platform at 4°C. The beads were then collected by centrifugation at 1500 rpm for 9 min in a Sorvall BP8 rotor at 4°C. The supernatant was discarded and the beads were resuspended in 500 ml Buffer H (20 mM Hepes-Cl, pH 7.5, 1 mM EDTA, 10% glycerol and protease inhibitors (Cytiva (#P8210)), followed by centrifugation at 1,500 rpm for 9 min in a Sorvall BP8 rotor at 4°C. This washing procedure was followed two more times, and then the beads were resuspended in 50 ml Buffer H, centrifuged once again, then resuspended in 5 ml buffer H and applied to a “C column” (Cytiva). The column was then connected to an ACTA FPLC and rinsed with buffer H + 1M NaCl until the OD_280_ no longer changed (about 3 column volumes). Elution was with buffer H + 300 mM KCl plus 0.2 mg/ml 3X-FLAG peptide, using three 1 column volume pulses (30 min). Proteins were visualized in 10% SDS PAGE, pooled, and concentrated to 2-2.5 μM using a 30 kDa cut-off filter to remove FLAG peptide. Proteins were aliquoted and stored at -80°C.

### DNAs

DNA oligonucleotide substrates were ordered from (Integrated DNA Technologies, Coralville, IA). Sequences used in this report are:

DNA Template Strand: TGT GGT AGG AAG TGA GAA TTG GAG AGT GTG GGT GAG GGT TGG GAA GTG GC

RNA Template Strand: rUrGrU rGrGrU rArGrG rArArG rUrGrA rGrArA rUrUrG rGrArG rArGrU rGrUrG rGrGrU rGrArG rGrGrU rUrGrG rGrArA rGrUrG rGrC

Primer: GTC TCG AGC CCA TCC TTC CAC TTC CCA ACC CTC ACC

Biotin Primer: GTC TCG AGC CCA TCC /TEGBiotin/ TTC CAC TTC CCA ACC CTC ACC

### Substrates

Oligo pair substrates were annealed in 100mM Tris-Cl pH 7.5, 1 mM EDTA, 100 mM KCl at a final concentration of 120 nM labeled primer with 180 nM unlabeled template strand to ensure saturation of the labeled primer. Two primary substrates were annealed, containing the primer and either the DNA or RNA template strand. Substrates were used at a final concentration of 1 nM in all reactions.

### Primer extension assay

Reactions were performed in reaction buffer (20 mM Tris-Acetate pH 7.5, 4% glycerol, 0.1 mM EDTA, 40 μg/mL BSA, 5 mM DTT, and 10 mM MgSO_4_). To initiate reactions, polymerase mixes containing the four dNTPs (100 μM final concentration each) and polymerase (20 nM final concentration, unless otherwise indicated) were added to substrate mixes containing either RNA or DNA template strand substrates, each primed with a 5’-^32^P end-labeled DNA oligo. Reactions were incubated at 30°C for the indicated length of time. Positive control lanes are cM-MuLV RT (New England Biolabs, Ipswich, MA) and negative control lanes are substrate mixes prior to polymerase addition. For reactions containing RFC-PCNA, a modified substrate was used in which the primer oligo has a TEGBiotin modification and the substrate was pre-bound with RPA to prevent loaded PCNA from sliding off the short substrate. For reactions containing RPA, substrate mixes with and without RPA (100 nM final concentration) were incubated for 5 min at 30°C to allow RPA to bind the ssDNA before reactions were initiated upon addition of polymerase. Reactions were terminated by adding an equal volume of 2x Stop buffer (1% SDS, 50 mM EDTA, 8% glycerol, .01% bromophenol blue, 83% formamide, 100 nM unlabeled primer oligo), boiled for 5 minutes, and analyzed in a 12% Urea PAGE gel. Gels were exposed to a phosphor imaging screen for 16-18 hrs then scanned with an Amersham Typhoon.

### RPA gel shift assay

Reactions were set up in the same buffer conditions as in the primer extension assay described above. 100 nM RPA was incubated with 1 nM DNA primed RNA template for 5 minutes at 30°C. One tenth volume of 10x loading dye (80% glycerol, 0.1% bromphenol blue) was added to each reaction, and samples were analyzed in a 10% native PAGE gel.

## Acknowledgement

This study was supported by the US National Institutes of Health grant GM148159 and the Howard Hughes Medical Institute (to M.E.O).

---

**Figure S1.**
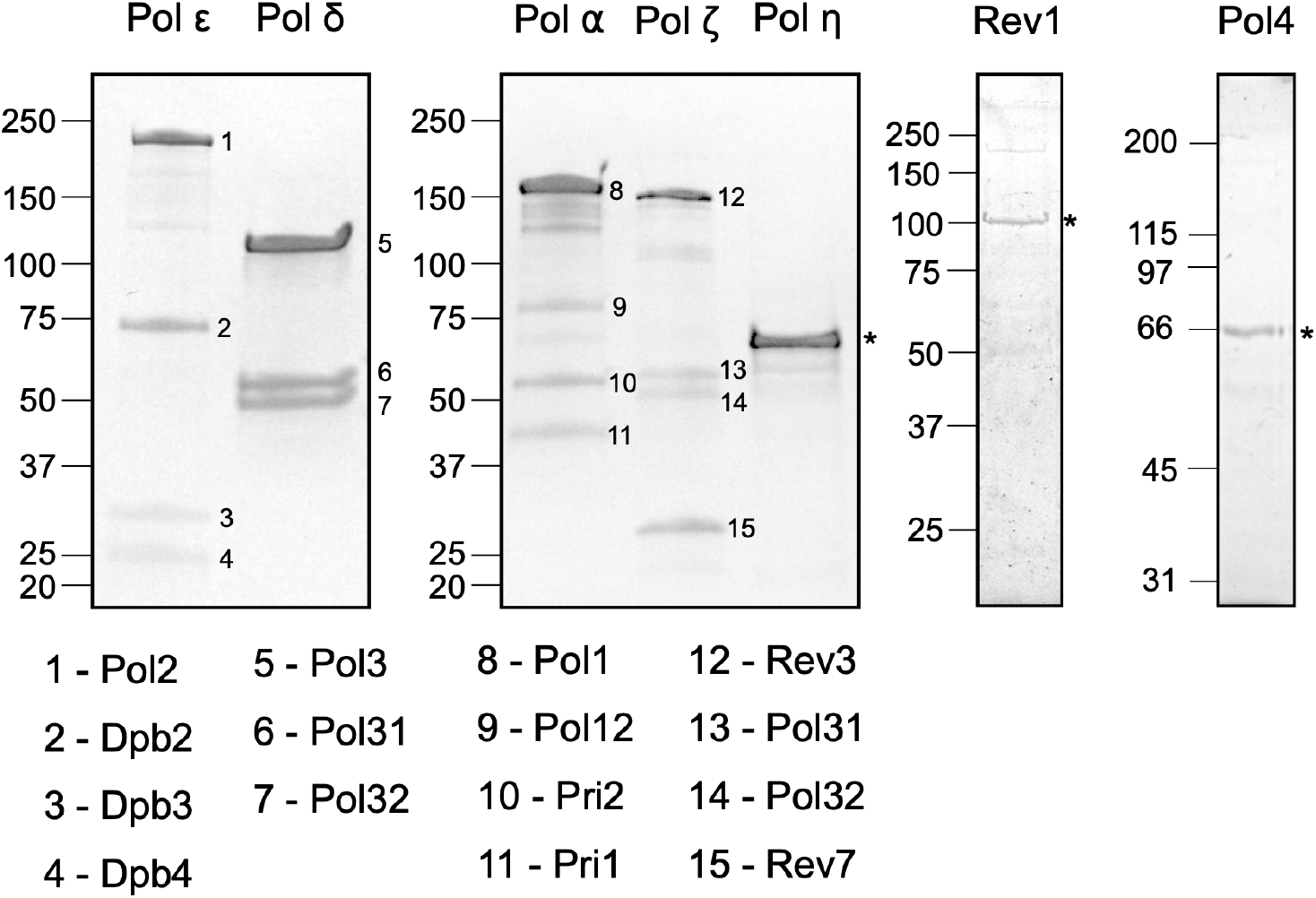
SDS-PAGE gels showing each of the purified yeast polymerases: Bands for individual proteins are indicated by either a number, for multi-subunit enzymes, or a * for single subunit polymerases.

**Figure S2.**
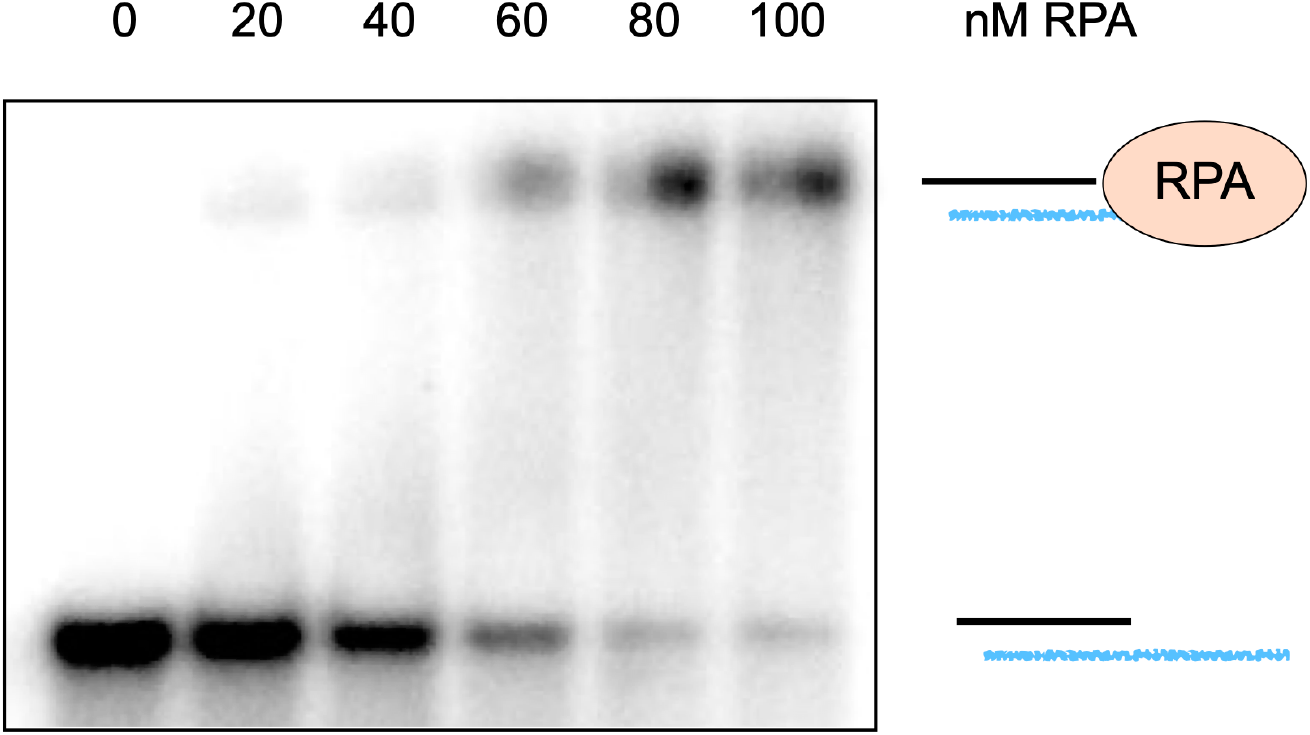
RPA binds the DNA primed RNA substrate: A gel shift assay after incubation of RPA with the DNA primed RNA substrate for 5 minutes at 30°C.

**Figure S3.**
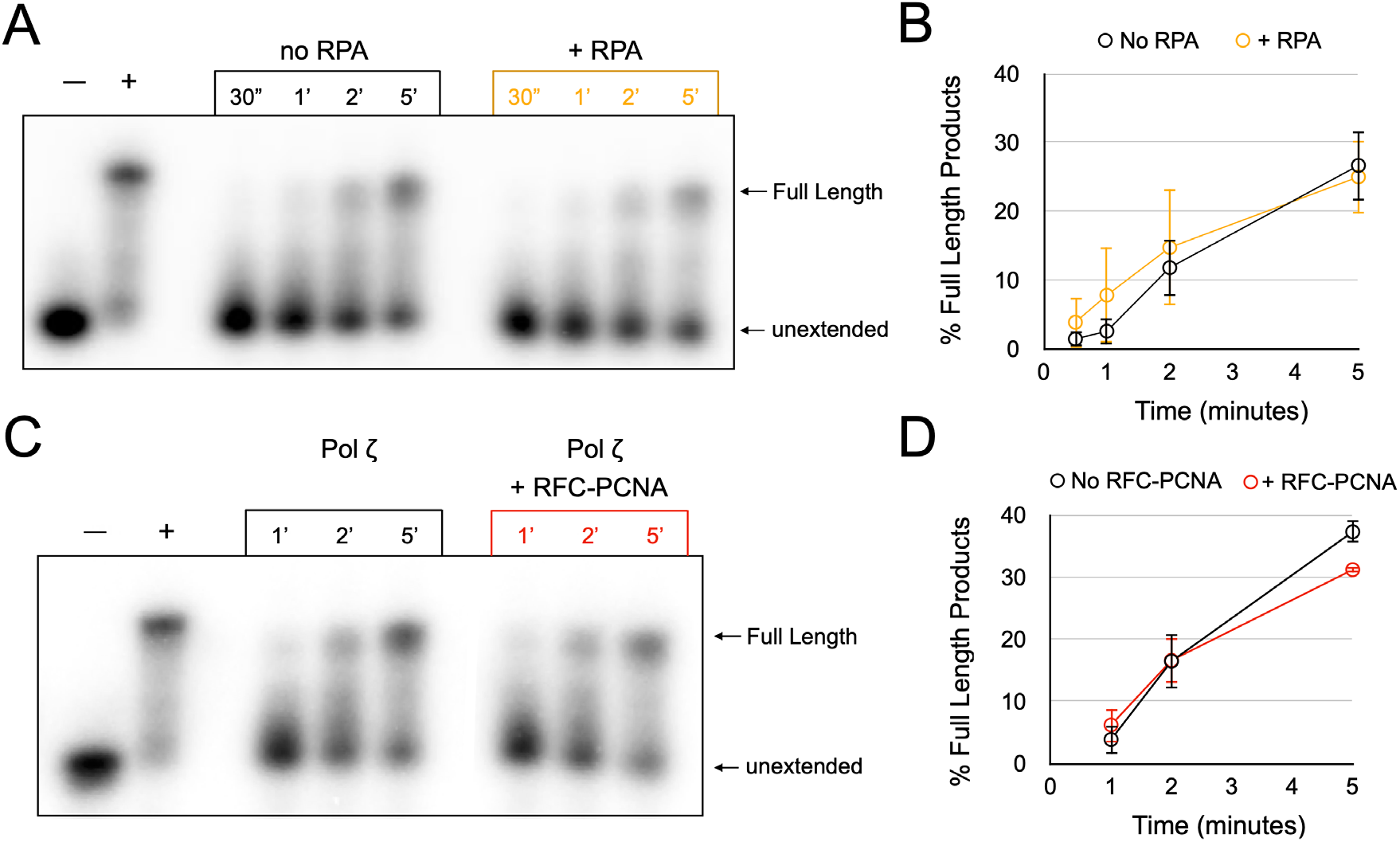
RPA and RFC/PCNA have no notable impact on Pol ζ RT activity: **(A)** shows primer extension RT assays comparing the presence and absence of RPA, and **(B)** is a quantitation of triplicate assays. Error bars represent +/-standard error of the mean. **(C)** shows primer extension RT assays comparing the presence and absence of RFC/PCNA. **(D)** is a quantitation of triplicate assays. Error bars represent +/-standard error of the mean.

**Figure S4.**
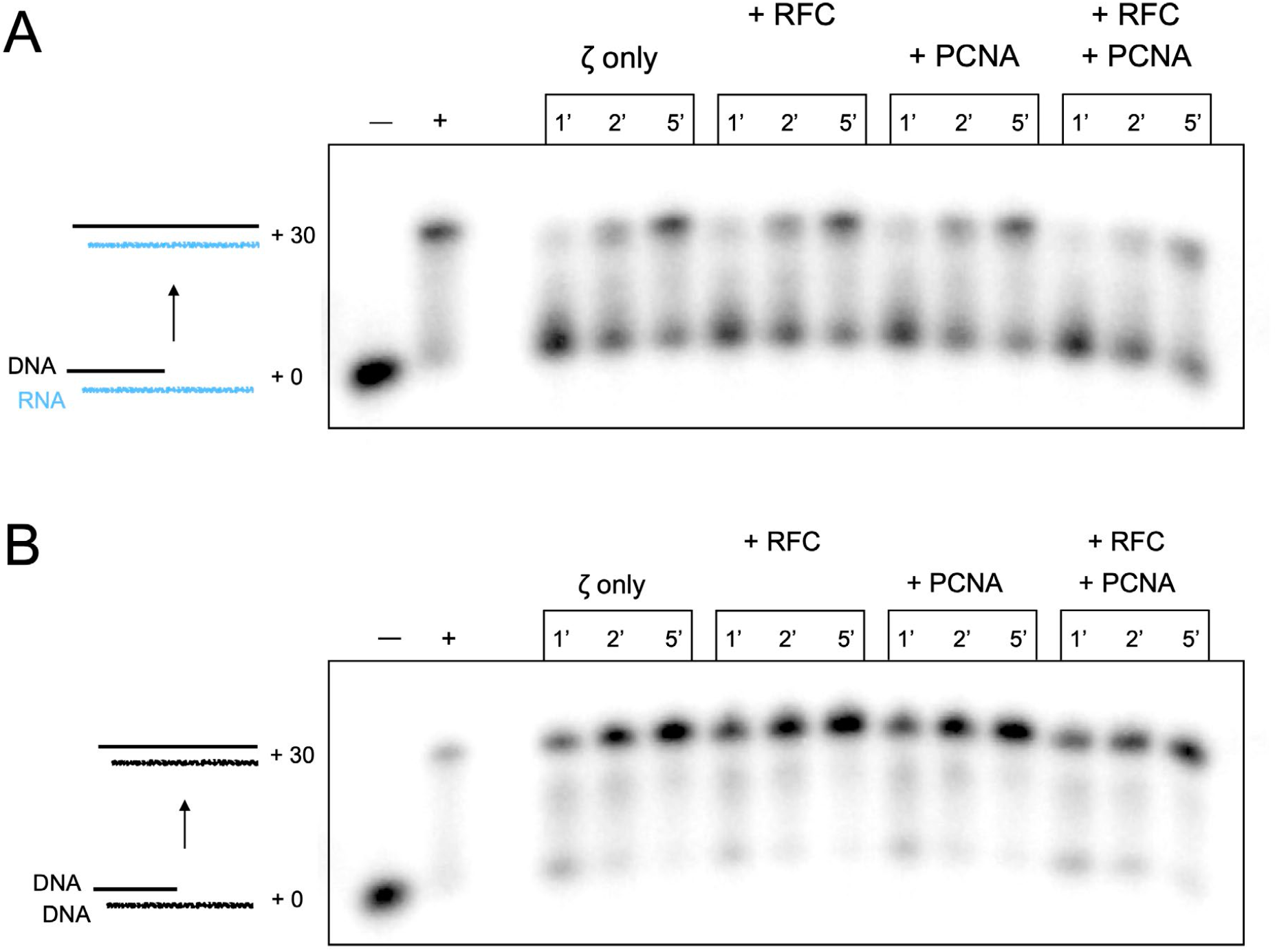
RFC +/-PCNA does not stimulate Pol ζ on either DNA or RNA templates: Extension assays with the noted additions of RFC and/ or PCNA using either RNA (A) or DNA (B) template strands.

